# Decoding Tumour-Specific Rewiring and Synthetic Lethality Through Genome-Scale Metabolic Models

**DOI:** 10.64898/2026.08.21.746174

**Authors:** Maziya Ibrahim, Rutuja Bhoite, Karthik Raman, Meiyappan Lakshmanan

## Abstract

Cancer cells rapidly rewire their metabolism, from efficient energy production toward anabolic processes, to sustain uncontrolled growth. Decoding such metabolic shifts is essential for uncovering novel therapeutic targets. To map systems-level metabolic changes across cancer types, we built context-specific genome-scale metabolic models for eight tissues (lung, thyroid, stomach, prostate, liver, kidney, colon, and breast) using gene expression data from The Cancer Genome Atlas (TCGA). Applying constraint-based modelling, we then identified differentially regulated pathways through flux enrichment analysis, revealing tissue-specific rewiring: branched chain amino acid metabolism was suppressed in breast cancer; sphingolipid metabolism was downregulated in colon, kidney, and thyroid but upregulated in breast. We further propose a model-driven pipeline to identify and characterise metabolic vulnerabilities. We first identify synthetic lethal reactions in normal tissues and their corresponding single lethal counterparts in cancers, thereby enabling the identification of metabolic “collateral lethal” reaction pairs for each cancer. Model-predicted collateral lethal gene pairs, including *CMPK1–AK* in colon, *ALDOA–PGD* in prostate, and *SLC25A26–UQCRB* in liver models, were supported through computational validation using DepMap data on gene essentiality. Subsequently, we show how to interpret metabolic rewiring in cancer tissues while accounting for any collateral lethal pairs. In summary, our results establish a systemic framework for decoding metabolic rewiring and synthetic lethal vulnerabilities in cancer.

**Author Summary:** Cancer cells alter their metabolism to support rapid growth and survival, but these metabolic changes can differ substantially between cancer types. Understanding which metabolic changes are shared across cancers and which are tissue-specific is necessary for identifying selective therapeutic opportunities. Here, we use genome-scale metabolic models to investigate how metabolism changes between normal and tumour states across eight tissues. By integrating tissue-specific gene expression with metabolic models, we generated context-specific metabolic models that capture the metabolic capabilities of individual tumour and adjacent-normal tissues. We found that cancer-associated metabolic rewiring is predominantly tissue-specific, with distinct changes in metabolic pathways and reaction usage across cancer types. Importantly, no single metabolic reaction was essential across all cancer tissues, highlighting the limited potential of universal metabolic targets. We also implement a collateral lethality framework to identify metabolic dependencies that arise specifically in tumour models. Some of the predicted tissue-specific collateral-lethal gene pairs were corroborated by independent cancer-dependency data from the DepMap resource. Finally, we used these predicted dependencies to examine the metabolic rewiring associated with the transition from normal to tumour states. Together, our findings demonstrate how genome-scale metabolic modelling can connect cancer-associated metabolic rewiring to tissue-specific dependencies and generate testable hypotheses for selective cancer targeting.

## 1. Introduction

Metabolic reprogramming, a hallmark of cancer, is well known to drive the rapid proliferation and survival of malignant cells, yet the specific metabolic rewiring that underpins this process across different organs remains a critical, unanswered question. Over the past few decades, research on cancer metabolism has revealed several aspects of this reprogramming. It is primarily manifested through three interconnected mechanisms: altered bioenergetics, enhanced biosynthesis, and maintenance of redox balance [1]. A seminal example of such reprogramming is the *Warburg effect*, described by Otto Warburg, in which cancer cells preferentially consume glucose and glutamine, and secrete lactate, even under normoxic conditions [2]. This shift not only supports energy production but also diverts glycolytic and TCA cycle intermediates toward the synthesis of macromolecules essential for rapid cell proliferation. Similarly, alterations in lipid metabolism, including increased *de novo* lipogenesis and the serine-glycine-one-carbon metabolism pathway, which supports nucleotide synthesis and redox balance, were also reported to be crucial for tumour proliferation [1]. Metabolic synthetic lethality linked to TCA-cycle mutations in fumarate hydratase, isocitrate dehydrogenase, and succinate dehydrogenase creates dependencies on other pathways, including NAD+ salvage and ROS handling. Rational combinations of inhibitors targeting glycolysis, oxidative phosphorylation, and lipid metabolism, such as *ACLY* and *FASN*, are necessary to overcome metabolic plasticity, which varies with patient-specific mutations, and to enable personalised treatment [1].

Recent advances in high-throughput omics technologies, combined with computational modelling, have significantly enhanced our ability to characterise and interpret metabolic alterations in cancer. One of the most powerful tools in this domain is the genome-scale metabolic model (GEM), which provides a comprehensive representation of cellular metabolism. Reconstructions such as Recon 2 [3] have laid the foundation for modelling human metabolism at a systems level. These models are typically analysed using constraint-based modelling approaches, such as flux balance analysis (FBA), which impose physicochemical constraints, including mass balance, thermodynamic feasibility, and reaction capacity, to predict feasible metabolic states that satisfy a defined objective, such as biomass production or ATP yield.

The application of genome-scale metabolic modelling to cancer research has yielded promising insights [4,5]. Since GEMs capture the complexity of interconnected metabolic pathways, they provide a rigorous framework for systematically analysing perturbations within the network. Some of the earliest attempts to integrate GEMs with cancer metabolic data included building 280 models from cancer and normal cell-line data and predicting drug targets, such as *MLYCD*, that selectively target cancer cells [6]. Lewis et al. used metabolic models integrated with transcriptomic and genomic data from 915 TCGA tumour samples to identify metabolic biomarkers distinguishing radiation-sensitive from radiation-resistant tumours, aided by statistical modelling-based multi-omics classifiers [7]. Similarly, a GEM of pancreatic ductal adenocarcinoma, incorporating patient-specific transcriptomics, has uncovered metabolic pathway divergences between malignant and normal tissues, leading to predictions of potential drug repurposing opportunities [8]. In another study, integrating gene expression profiles from 1,156 breast cancer and normal samples with human GEMs enabled the identification of breast cancer-specific metabolic signatures associated with androgen and estrogen metabolism, fatty acid oxidation and the TCA cycle [9]. Furthermore, a pan-cancer study [4] reconstructed genome-scale metabolic models for 917 primary tumours across 13 cancer types, revealing that core reactions in tumour networks largely overlap with normal housekeeping cellular functions. This network-centric approach demonstrated that contextual metabolic variations are primarily dependent on cancer type, indicating that tissue-specific metabolic adaptations dominate over universal cancer rewiring. While these studies focused on analysing the metabolic rewiring of individual cancers and comparing it across multiple cancers, they have not compared the metabolism of pan-cancers with that of the corresponding normal or tumour-adjacent normal tissues.

In the current study, we present a systems-level computational framework for identifying tissue-specific metabolic vulnerabilities in cancer, capitalising on the structure of GEMs and the principles of constraint-based metabolic modelling. While GEMs have been used to analyse cancer metabolism in individual tissues, they have not been used extensively to characterise shifts in metabolic pathway fluxes from normal to a cancer state. Here, we adopt a systems-level strategy that uses GEMs to distinguish tumour-specific metabolic dependencies from those of adjacent-normal tissue and further identifies collateral lethal gene targets that can be exploited to preferentially impair cancer cell survival (Fig.1). It should be noted that while we demonstrated this strategy with eight cancer tissues, it can be extended to other cancer types with matched normal gene expression data, underscoring the utility of the GEM-based collateral lethality screening approach.

**Fig. 1.**
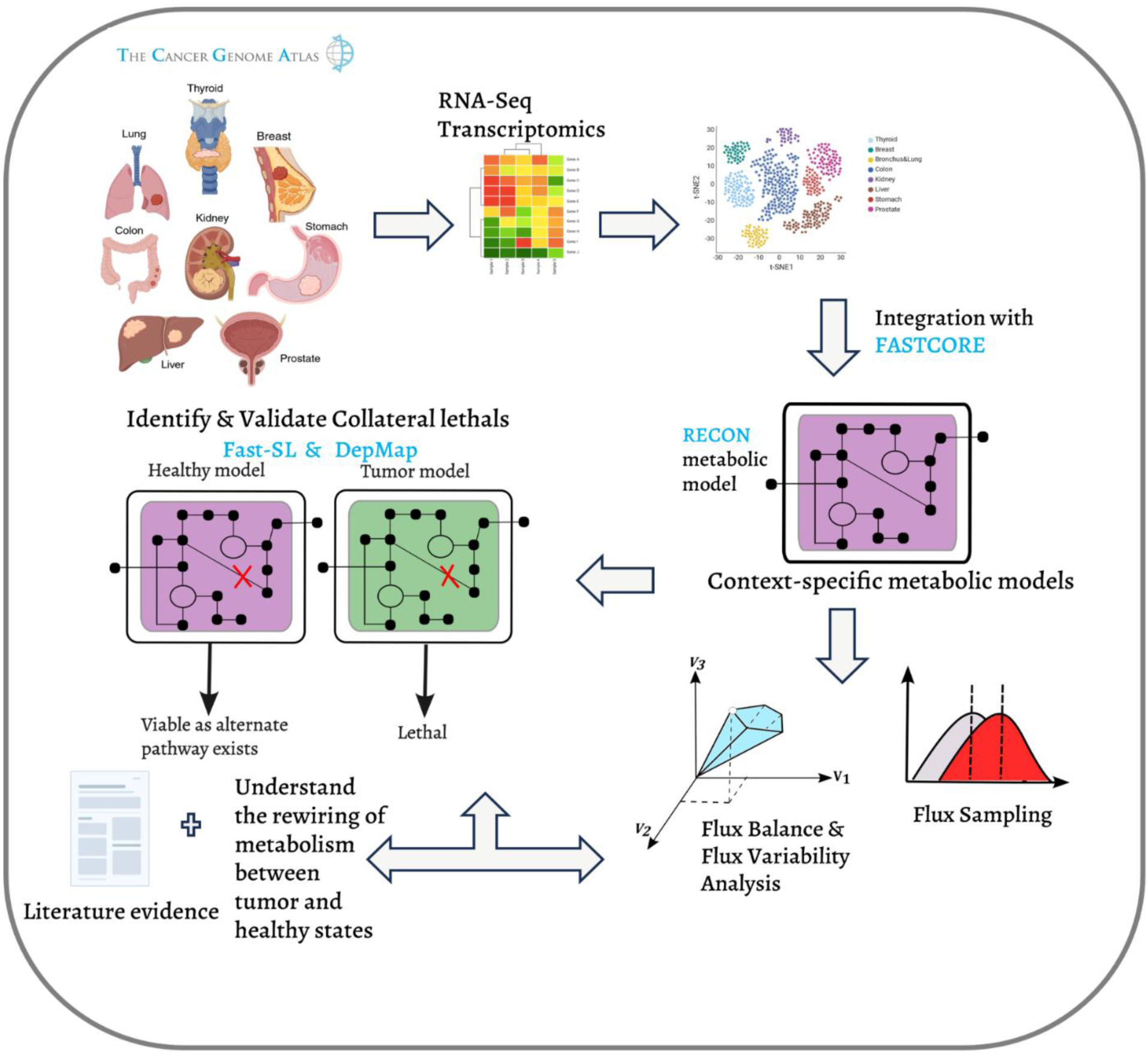
Overview of the metabolic modelling approach used in the study.

## 2. Results

### 2.1 Reaction profiles reveal that tissue-specific divergence outweighs pan-cancer metabolic similarities

To explore the genome-wide metabolic landscapes in both cancer and normal conditions across multiple tissues, we reconstructed 480 context-specific genome-scale metabolic networks. Using the gene expression data extracted from the TCGA database for eight tissues, namely breast, bronchus and lung, colon, kidney, liver, prostate, stomach, and thyroid, we established the GEMs representing both normal and malignant tissues using the FASTCORE [10] algorithm. We used t-distributed stochastic neighbour embedding (t-SNE) [11], an unsupervised nonlinear dimensionality reduction technique, to cluster and visualise metabolic model data by tissue type and condition (i.e., normal or tumour), as well as gene expression data from the same tissues. We observe that the models cluster well by tissue condition (Fig. 2D), but there are mixed clusters by tissue type (Fig. 2C), as seen in models representing stomach and colon tissues. Gene expression appears more similar to the tissue of origin than to that of other cancer tissues (Fig. 2A & Fig. 2B). This suggests that there are fewer pan-cancer signatures and more tissue-specific signatures in the cancer state. The concordance between gene expression data and model-derived profiles provides evidence that the reconstructed models capture biologically relevant signals.

**Fig. 2.**
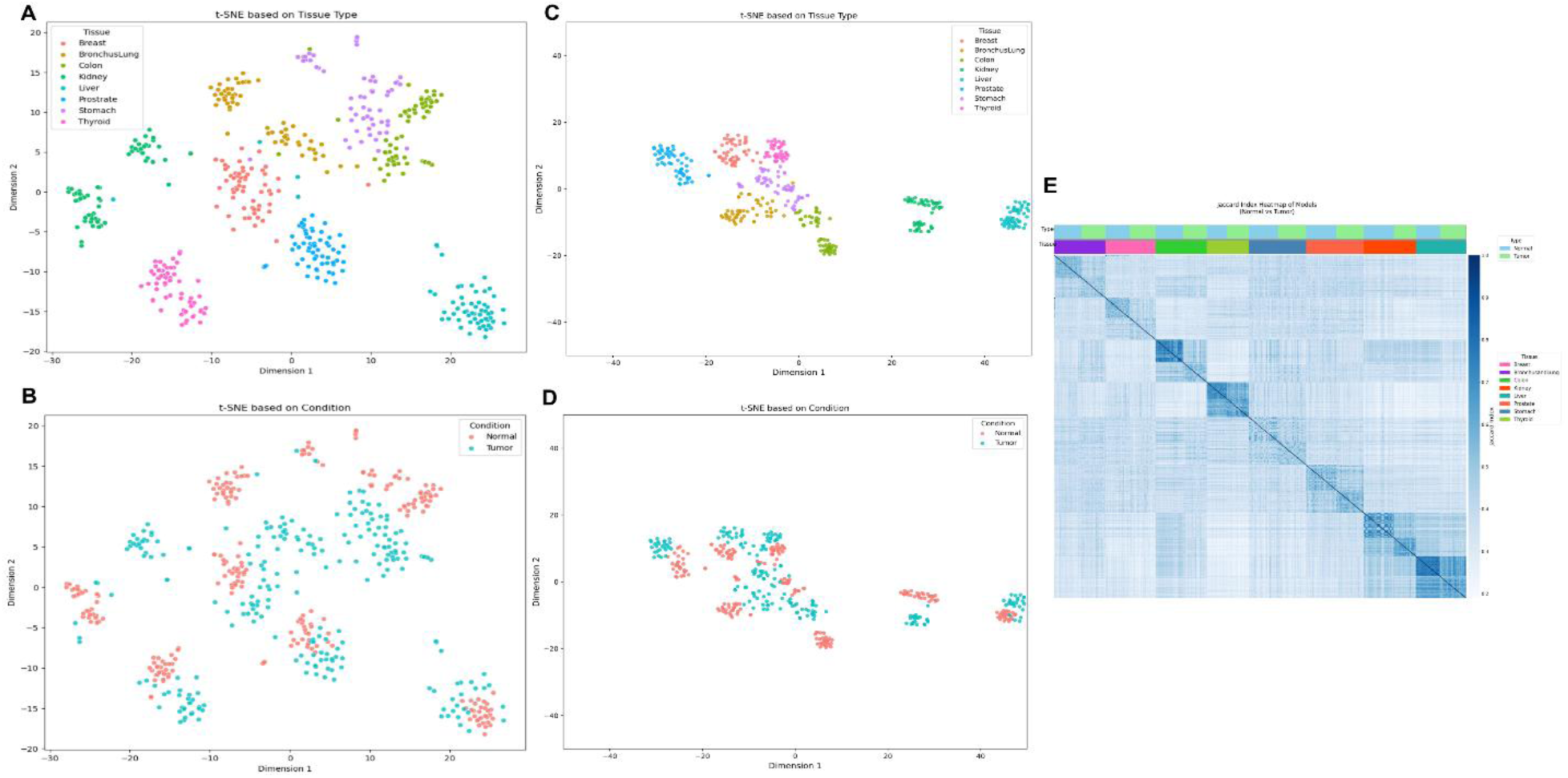
t-SNE clustering based on gene expression profiles and context-specific metabolic models. **2A–2B:** Clustering using gene expression data from eight tissue types. **2A** is coloured by tissue type, while **2B** is coloured by condition (normal or tumour). **2C–2D:** Clustering of context-specific metabolic models based on tissue type and condition, respectively. **2E:** Heatmap showing pairwise Jaccard similarity indices among all models, grouped by tumour status and tissue type.

With pairwise Jaccard indexes computed across all models and tissues, we observe that certain tissues, such as the colon and kidney, exhibit a clear distinction between normal and tumour conditions, with Jaccard similarities ranging from 0.2 to 0.4. Whereas certain tissues, such as the liver and thyroid, have models with greater similarity between normal and tumour conditions. Liver and kidney tissue models appear to share similar sets of reactions.

Model statistics on the number of reactions in each tissue type show that most tissues, including the lung, colon, kidney, liver, and stomach, exhibit significantly different reaction counts between normal and cancer conditions (Fig. 3A and Supplementary Table S1). Kidney tissue shows the most distinct differences between normal and tumour states. To gain deeper insight into the functional metabolic capabilities of these models, we computed core and accessory reactions (Fig. 3B). A reaction was defined as ‘core’ if it is present in at least 80% of the cancer metabolic models in a particular tissue. Accessory reactions are those that are shared between two or more tissue types, while unique reactions are found only in one tissue.

**Fig. 3.**
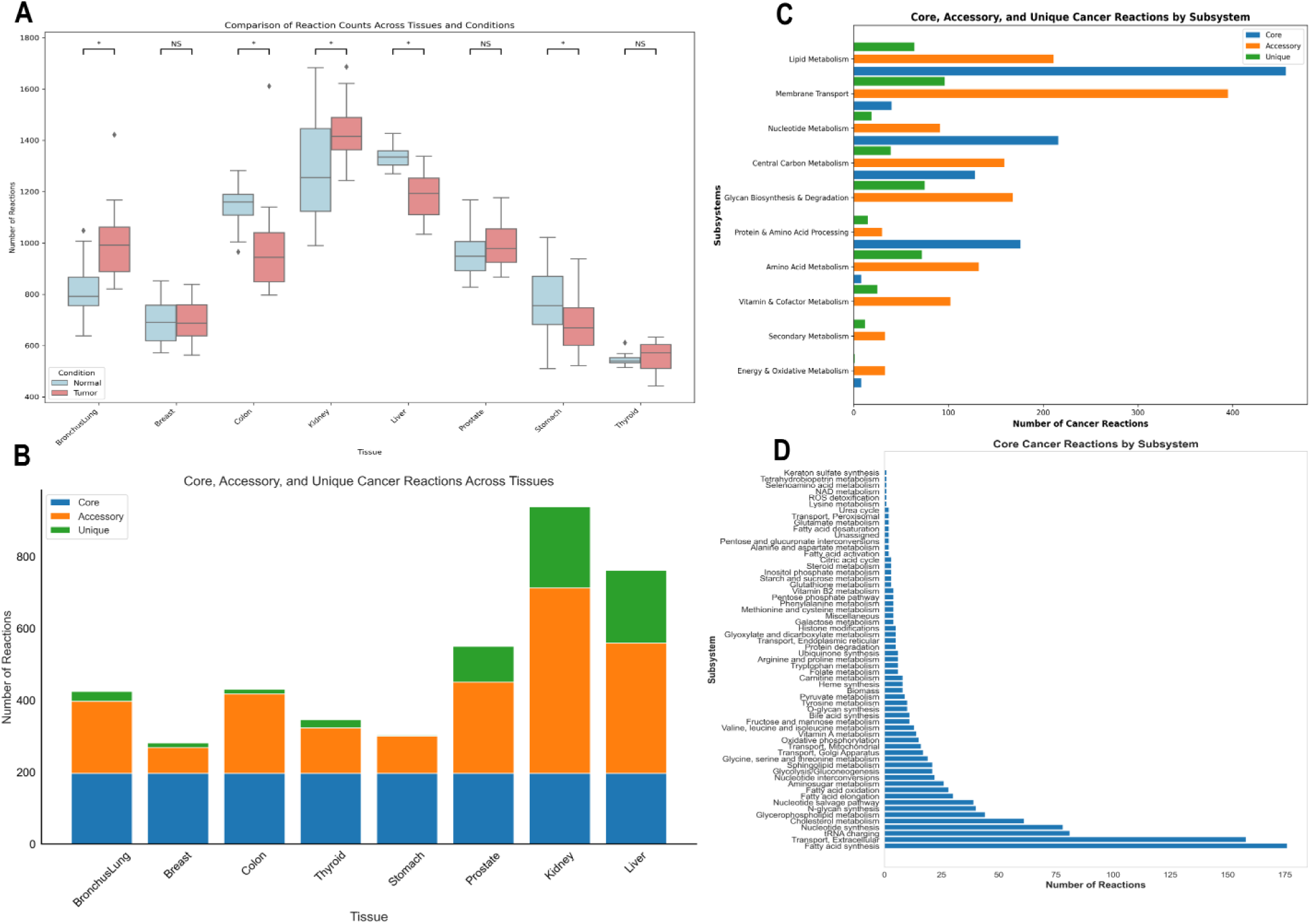
Metabolic diversity across cancers from different tissues, based on core, accessory, and unique reaction and subsystem profiles. **3A:** Distribution of reaction counts in context-specific metabolic models across eight tissue types under normal and tumour conditions. Statistical significance between conditions is indicated for each tissue type (*p* < 0.05; NS, not significant). **3B:** The number of core, accessory, and unique cancer reactions identified across different tissues. **3C:** Distribution of core, accessory, and unique cancer reactions across major metabolic subsystems, highlighting subsystem-specific metabolic diversity in cancer. **3D:** Breakdown of core cancer reactions by detailed metabolic subsystems, illustrating the dominant pathways conserved across cancer tissues.

Prostate, kidney, and liver tissue models exhibit a higher number of accessory and unique reactions compared to other cancer types (Fig. 3C). Most accessory reactions are associated with transport processes, suggesting context-dependent modulation of nutrient uptake and exchange between cellular compartments (such as mitochondria and the extracellular space) to meet the specific metabolic demands of different cancer types. In contrast, most core reactions conserved across cancer models are involved in nucleotide, lipid, and amino acid metabolism. A detailed examination of the pathways linked to these core reactions reveals that many are integral to fundamental metabolic processes such as fatty acid synthesis, cholesterol metabolism, the nucleotide salvage pathway, and glycolysis (Fig.3D). Reactions involved in keratan sulphate synthesis and lysine metabolism represent a minority among the core reactions identified in cancer models. Overall, the tissue models exhibit a spectrum of metabolic plasticity, with some cancers, such as those of the stomach and breast, having more conserved pathways, while others, as discussed above, have more specific metabolic capabilities, particularly in the kidney and liver.

### 2.2 Cancer metabolic networks exhibit universal lactate upregulation but tissue-dependent amino acid changes

Constraint-based models of metabolic networks are typically mathematically an underdetermined linear system, because they contain more reactions than metabolites. Therefore, the solutions to this system do not consist of unique flux rates for each reaction, but rather a space for possible flux rates. By uniformly sampling this space, the distribution of sampled feasible fluxes can be obtained and statistically compared between normal and malignant states to identify metabolic shifts. Fig. 4 shows the reactions whose fluxes are up- or downregulated in each cancer tissue. These differentially active reactions were identified as mentioned in section 4.3. We observe that lactate flux is universally upregulated in all tissues, consistent with the increased glycolytic/lactate-producing phenotype characteristic of the Warburg effect.

**Fig. 4.**
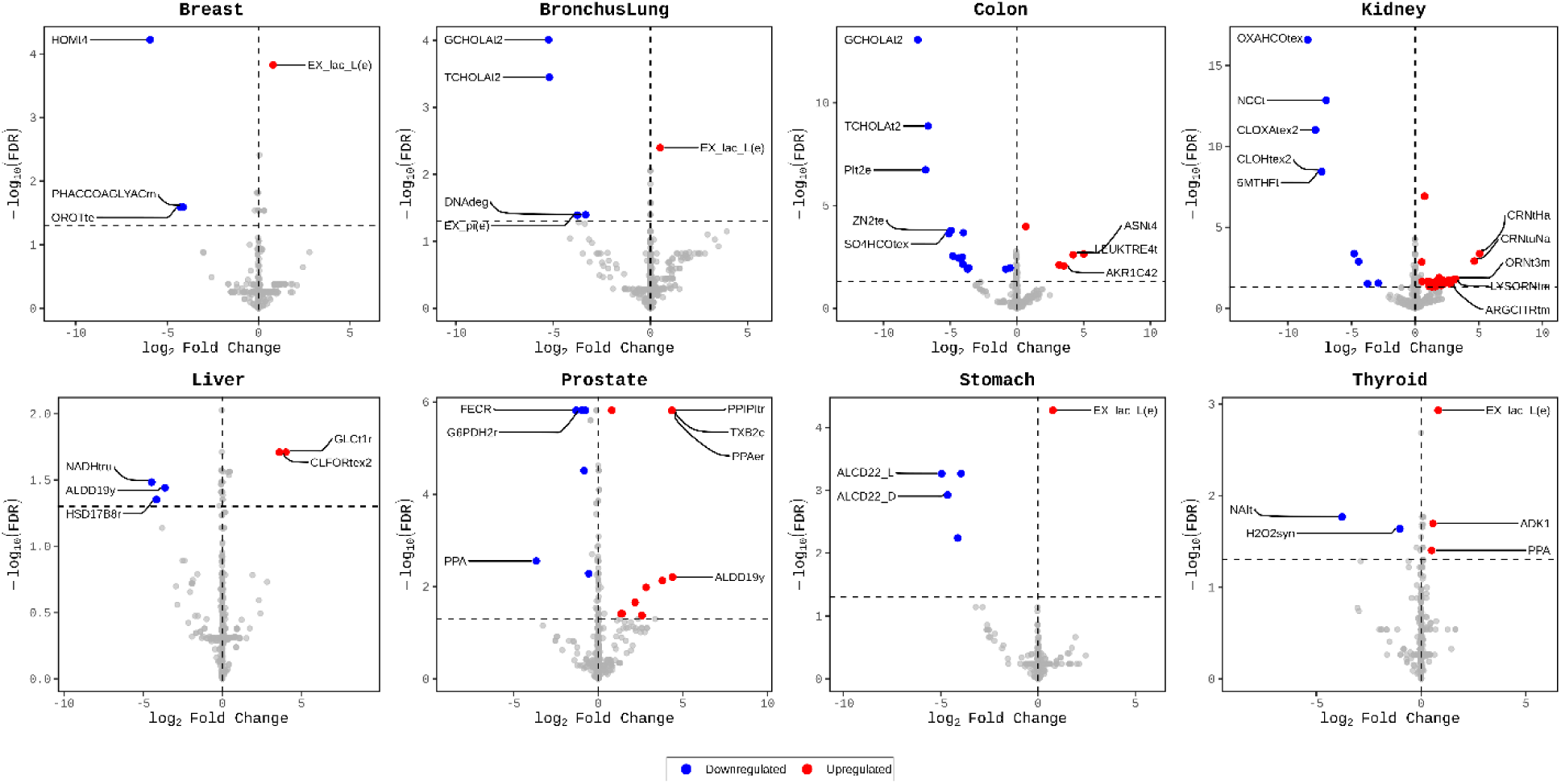
Differentially expressed fluxes of reactions in cancer tissue models. Reactions with up-regulated and down-regulated fluxes obtained from flux sampling in cancer models compared to normal tissue models.

We perform pathway enrichment analysis using the complete set of reactions and pathways from the Recon model. The analysis reveals that branched-chain amino acid (BCAA) metabolism is negatively enriched in breast cancer models, while keratin sulfate metabolism is specifically enriched in kidney models. Sphingolipid metabolism is downregulated in colon, kidney, and thyroid tissues, but upregulated in breast tissue. Tyrosine metabolism shows negative enrichment in the kidney, while it is positively enriched in prostate models. Pentose phosphate pathway and pyruvate metabolism are downregulated in prostate and stomach tissues, respectively. (Fig. 5).

**Fig. 5.**
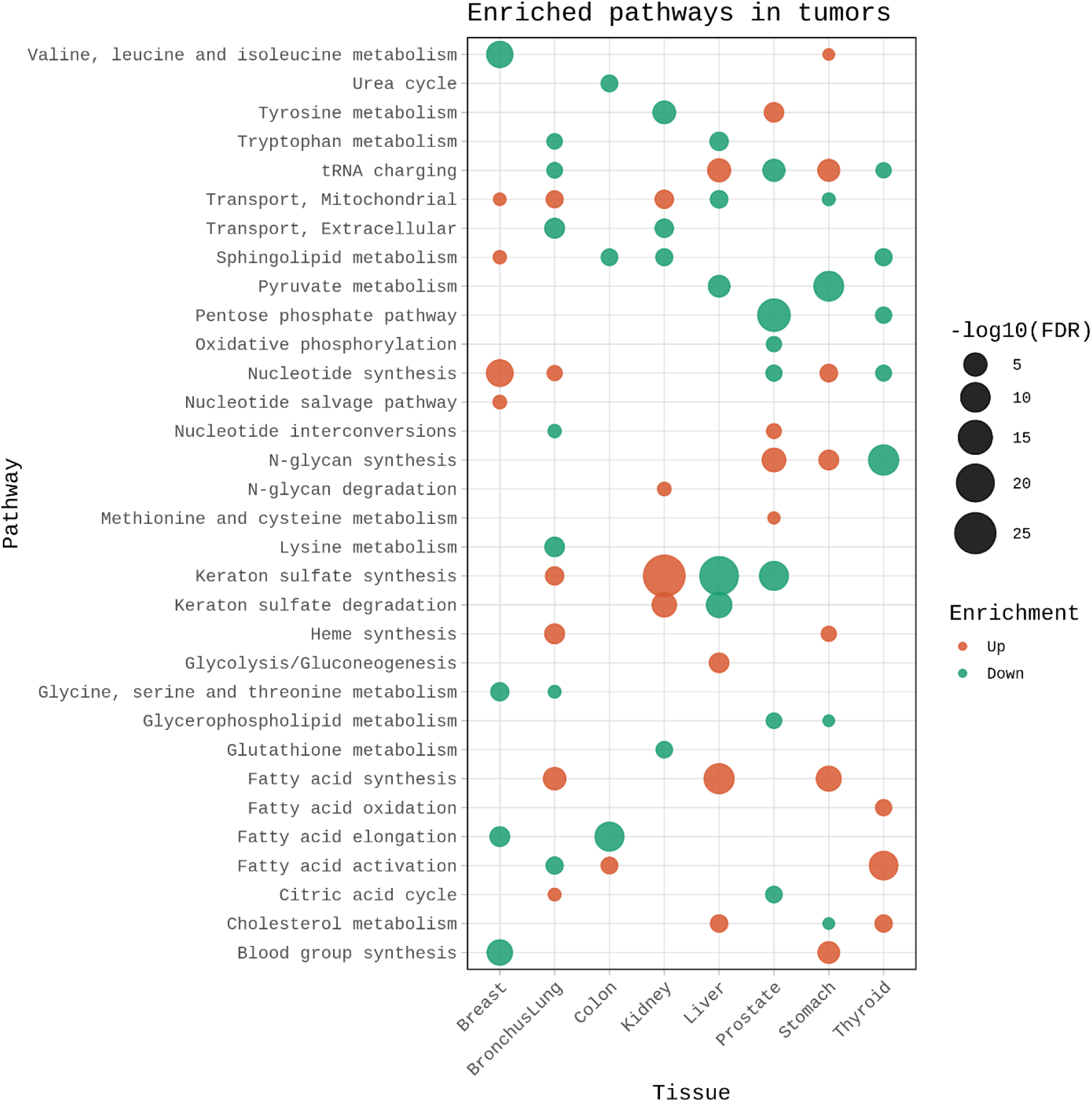
Enrichment of pathways in cancer models based on flux sampling

### 2.3 Essential reaction profiling highlights the absence of a universal pan-cancer target

Essential reactions, also termed ‘single lethal’ reactions across cancer tissue models, were identified using Fast-SL [12] (see Methods). As shown in Supplementary Fig. S1A, prostate cancer models harboured the largest number of essential reactions, followed by breast and kidney models, while lung models exhibited the fewest. Pairwise overlap analysis revealed that thyroid and prostate models shared the greatest number of essential reactions (*n* = 8), predominantly involving the pentose phosphate pathway, glycolysis, and transport reactions. Lung and colon models shared seven essential reactions, primarily from glycerophospholipid metabolism. Notably, no single reaction was found to be essential across all eight tissue types, suggesting the absence of a pan-cancer metabolic core at the reaction level. Tissue-specific essentiality was also evident at the pathway level (Supplementary Fig. S1B): breast cancer models were enriched for essential reactions in the TCA cycle, while kidney models showed pronounced dependencies on arginine, proline, and branched-chain amino acid (BCAA) metabolism.

### 2.4 DepMap validation of collateral lethality identifies tissue-specific vulnerabilities

Two or more biochemical pathways can exist for the same essential cellular processes. Any homozygous passenger deletions can affect one pathway while leaving the other undisturbed, thereby rendering the affected pathway essential without compromising the viability of the cancer cell. This offers an opportunity to design selective inhibitors targeting enzymes in the non-deleted pathway, thereby resulting in tumour cell death while leaving normal cells intact. This concept has been previously defined as ‘collateral lethality’ ^11^. In the context of metabolic modelling, we have defined collateral lethality as a gene or reaction that is single lethal in cancer-specific tissue models but double-lethal in normal tissue metabolic models. As outlined in the methods section, we used Fast-SL [12] to identify and DepMap to validate the collateral lethal specific to each tissue type and across all tissues. Of all the collateral lethal reactions predicted for each tissue, we find no overlap across cancers, suggesting tissue-specific metabolic rewiring.

Using information on gene essentiality and mutations in cancer cell lines from DepMap, we verified our predictions of collateral lethal reactions. After mapping them to their corresponding genes, we identified 52 such lethal gene pairs, comprising 17 distinct genes across breast, colon, liver, and prostate tissues (Supplementary Table S2). Upon further comparison with data from OGEE, we find that three genes, *CMPK1* from the colon and *ALDOA* from the prostate, are included in the pan-cancer essential list in OGEE, and *SLC25A26* is listed as essential for liver tissue.

In colon tissue, we identified cytidine monophosphate kinase (*CMPK1*) and Adenylate kinase (*AK*) as a collateral-lethal gene pair. For prostate tissue, the predicted pair was aldolase and phosphogluconate dehydrogenase. In liver tissue, we predicted a potential collateral lethal association between the solute carrier gene *SLC25A2c* and ubiquinol-cytochrome reductase.

### 2.5 Mapping metabolic rewiring from normal to tumor state using in silico knockouts and FBA

To characterise the broader metabolic context in which the identified collateral lethalities arise, we investigated subsystem-level metabolic rewiring between normal and tumor states. We simulated the targeted deletion (see Methods) of the normal-tissue counterparts for our predicted pairs: adenylate kinase reactions in colon normal models, ubiquinol-cytochrome reductase reactions in liver normal models, and phosphogluconate dehydrogenase reactions in prostate normal models. Using FBA to derive reaction activity states, we then compared reactions that were exclusively active in either the cancer or normal tissue models. This allowed us to map the system-wide metabolic shifts associated with the malignant state. Fig. 6A and 6B illustrate the distribution of these differentially active reactions across metabolic pathways in colon and liver tissues, respectively (see Supplementary Fig. S2 for prostate tissue).

**Fig. 6.**
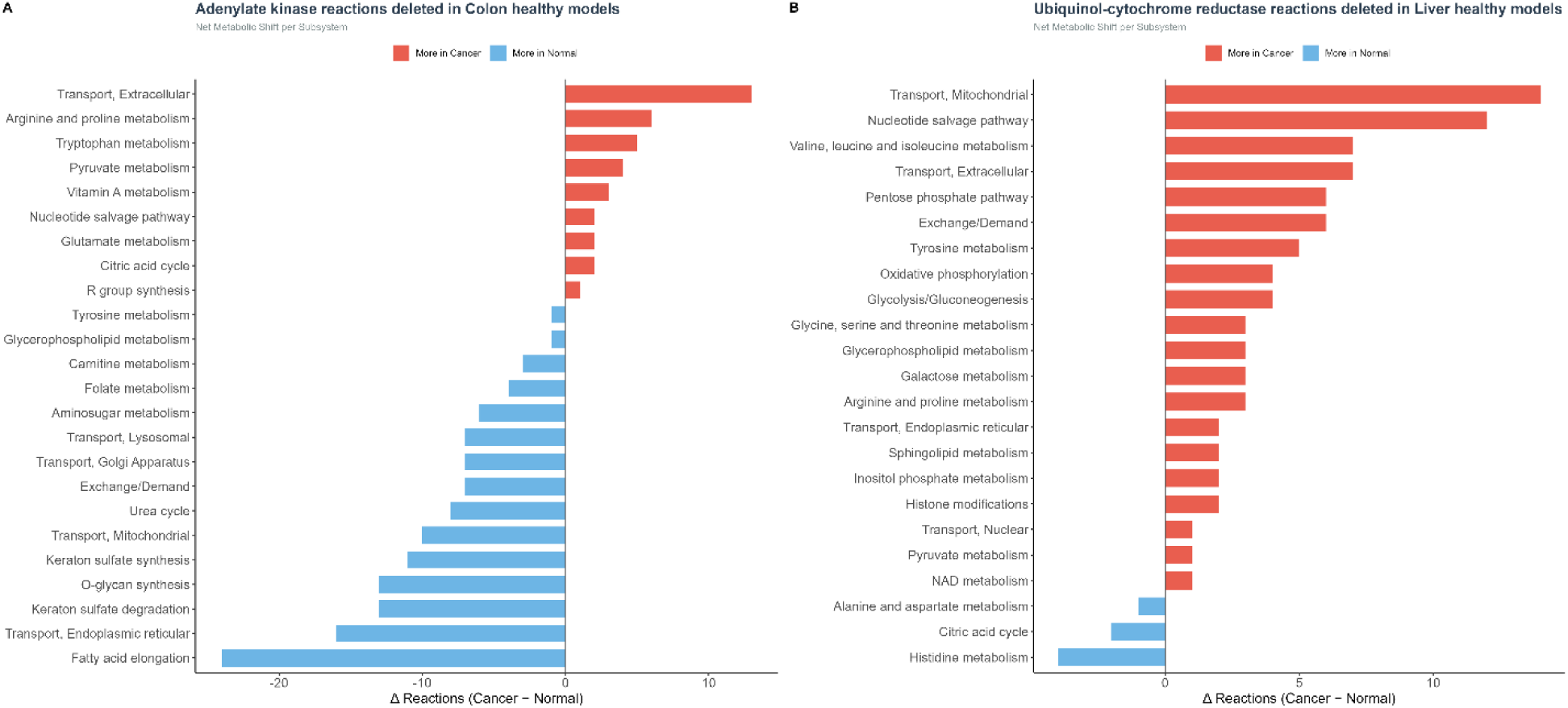
Metabolic rewiring in colon and liver tissue models. **6A:** Adenylate kinase reactions were deleted from normal colon tissue models, and the resulting metabolic rewiring was evaluated by comparing the number of reactions within subsystems that showed higher activity in cancer versus normal models. **6B:** Ubiquinol-cytochrome reductase reactions were deleted from normal liver tissue models, and the resulting metabolic rewiring was evaluated by comparing the number of reactions within subsystems that showed higher activity in cancer versus normal models.

In colon tissue models, reactions associated with nucleotide salvage, vitamin A metabolism, and amino acid metabolism pathways, such as those for arginine, proline, and tryptophan, were preferentially active in cancer models. In liver tissue models, mitochondrial transport, fatty acid oxidation, pentose phosphate pathway activity, and glycolysis/gluconeogenesis were enriched in cancer models. In prostate tissue models, tyrosine and glutathione metabolism, along with histidine metabolism, were preferentially active in cancer models.

## 3. Discussion

The use of genome-scale metabolic models enables a systems-level framework for integrating omics data, offering a high-level perspective compared to gene correlations from traditional transcriptomics data analysis. Metabolic diversity was found to be more tissue-specific; for instance, kidney tissue models were enriched in sphingolipid and tyrosine metabolism. This observation is highly consistent with previous pan-cancer metabolic network analyses, despite the differences in input data and context-specific model reconstruction algorithms [4,10,14]. Collectively, it is evident that the metabolic rewiring in cancer is predominantly tissue-specific rather than a universal pan-cancer metabolic core. Importantly, our observed trends were independent of the selected reference GEM, with Recon3D-derived models displaying equivalent clustering behaviour (Fig. S3).

Using flux sampling, we observe altered metabolism in cancer models. We demonstrate substantial tissue-specific metabolic rewiring across cancer models, with dysregulation of branched-chain amino acid (BCAA) metabolism in breast tissue models and altered tyrosine metabolism in kidney and prostate cancer models. These findings concur with recent advances reported in the literature on cancer research. Negative enrichment of branched-chain amino acid (BCAA) metabolic fluxes observed in breast cancer models may reflect suppression of oxidative BCAA catabolism and rerouting of BCAAs toward anabolic signalling pathways that support tumour growth. Consistent with this, Peng et al [15]. highlighted that BCAA metabolism is extensively reprogrammed in cancer, in a tissue-specific manner, in which reduced BCAA catabolism can enhance oncogenic signalling, such as mTOR activation, and promote tumour progression. Sphingolipid metabolism showed negative enrichment in colon, kidney, and thyroid cancer models, but positive enrichment in breast cancer models, indicating substantial tissue-specific rewiring of lipid metabolic programs. Given the functional heterogeneity of sphingolipid intermediates, including the opposing roles of pro-apoptotic ceramides and pro-survival sphingosine-1-phosphate, reduced pathway-level flux enrichment may reflect selective remodelling of specific sphingolipid branches rather than a uniform suppression of oncogenic sphingolipid signalling, consistent with previous reports of context-dependent sphingolipid metabolism in cancer [16]. Pyruvate metabolism was negatively enriched in stomach cancer models, potentially reflecting the altered routing of pyruvate away from oxidative mitochondrial metabolism and toward alternative metabolic programs characteristic of tumour metabolic plasticity. Such rewiring is consistent with studies showing that oncogenic signalling and microenvironmental adaptation dynamically regulate pyruvate utilisation and mitochondrial coupling in cancer cells [17].

We have identified novel candidate collateral lethal gene pairs across different cancers. These pairs include cytidine monophosphate kinase (*CMPK1*) and Adenylate kinase (*AK*) in colon tissue, aldolase and phosphogluconate dehydrogenase in prostate tissue, and *SLC25A26* and ubiquinol-cytochrome reductase in liver. While direct evidence for synthetic lethality or essentiality in cancer tissues remains limited, existing studies support links between these individual genes and cancer phenotypes. Adenylate kinases maintain adenine nucleotide homeostasis and regulate cell cycle control through the *AK–AMP–AMPK* axis [18]. In colorectal cancer, *AK6* is overexpressed and promotes migration and invasion by reinforcing glycolysis via interaction with lactate dehydrogenase A, whereas reduced AK5 expression is associated with enhanced proliferation, invasion, and altered *AMPK/mTOR* signalling, consistent with a tumour-suppressive role [19–21].

*CMPK1* activates the chemotherapeutic agent 5-fluorouracil by phosphorylating *FUMP*, and its miR-130b–mediated downregulation confers 5-FU resistance in gastric cancer [22]. In contrast, nuclear *CMPK1* promotes tumour growth in triple-negative breast cancer, indicating context-dependent functions [23]. Given the widespread use of pyrimidine antimetabolites in cancer therapy [24], the role of *CMPK1* in colorectal cancer warrants further investigation.

*SLC25A26* encodes the mitochondrial SAM transporter required for mitochondrial methylation. Its downregulation in hepatocellular carcinoma is associated with reduced expression of senescence markers, consistent with a tumour-suppressive role [25]. Reactivation of *SLC25A26* induces senescence via *AMPK/mTOR* and telomerase regulation, whereas impaired mitochondrial *SAM* import enhances oxidative phosphorylation, redox buffering, and chemoresistance, thereby promoting tumour survival [26,27]. These findings highlight *SLC25A26* as a potential therapeutic target.

Ubiquinol cytochrome reductase subunits have been implicated in the development of hepatocellular carcinoma. *UQCRH* and *UQCRB* are overexpressed in HCC and are associated with poor prognosis and hypoxia-driven angiogenesis. Pharmacological inhibition of *UQCRB* suppresses invasion and ROS production [28,29]. *UQCRB* upregulation has also been reported in colorectal cancer, whereas direct roles for *UQCRC2* and *UQCRFS1* remain unestablished [30]. *ALDOA* and *PGD* are key metabolic enzymes in glycolysis and the pentose phosphate pathway, respectively. Although our model simulations predicted an overall downregulation of pentose phosphate pathway flux in prostate cancer, the identification of PGD as a collateral lethal target aligns with literature showing targeted upregulation by androgen receptors signalling upregulates both genes, promoting a growth-amplifying feedback loop, while *ALDOA* overexpression correlates with poor prognosis and is pharmacologically targetable [31,32]. *PGD* also exhibits synthetic lethality in oxidative phosphorylation–deficient tumours [33]. While direct evidence of *ALDOA* and *PGD* in prostate cancer is limited, mechanistic parallels in other cancers [34] and pathway interdependencies seem to support this hypothesis.

Using these collateral lethal targets, we have synthetically uncovered rewiring of metabolic pathways in tumor tissues. Across colon, liver, and prostate cancer models, a consistent pattern of increased nucleotide salvage and amino acid metabolic pathway activity emerged following gene B deletion (as described in sections 4.4 & 4.5) in normal tissue models, reflecting the elevated biosynthetic and proliferative demands of the cancer state. In colon cancer, this is supported by evidence that *MYC*-driven upregulation of pyrimidine synthesis genes, including *CAD*, *UMPS*, and *CTPS*, is required for cancer cell growth, and that *MYC* similarly induces tryptophan transporters *SLC7A5* and *SLC1A5* to sustain kynurenine pathway flux, which functions as an oncometabolite in colon cancer [35]. The enrichment of retinol metabolism-associated genes observed in colorectal cancer organoids further corroborates the aberrant activation of vitamin A metabolic reactions predicted in our colon cancer models [36]. In liver cancer, the coordinated upregulation of glycolysis observed in our HCC models is consistent with the well-documented Warburg effect, in which increased glycolytic flux is coupled with PPP activation to meet biosynthetic and redox demands [37]. The predicted importance of mitochondrial transport in HCC is further supported by evidence that *SLC25A20* downregulation, which impairs acylcarnitine transport into mitochondria, promotes HCC growth and metastasis, suggesting that mitochondrial metabolite transport is a key metabolic axis in liver cancer, regardless of the specific transporter involved [38].

We recognise that, by design, genome-scale metabolic models represent enzymatic reaction networks and do not explicitly capture upstream regulatory processes, such as transcriptional programs and signalling cascades, that regulate metabolism. Consequently, the initial list of computationally predicted lethal reactions is large, and stringent biological filtering is necessary to distinguish credible candidates from false positives, a challenge inherent to all GEM-based lethality analyses. The three highlighted collateral-lethal gene pairs are supported by independent literature on their metabolic roles in cancer and represent prioritised candidates warranting further experimental validation. Taken together, our results provide a systematic framework for identifying collateral lethal vulnerabilities and lay the groundwork for context-specific therapeutic strategies in cancer.

Overall, our results highlight nucleotide salvage, amino acid metabolism, and mitochondrial transport as critical nodes in cancer metabolism, prioritise tissue-specific metabolic dependencies and candidate targets for experimental validation. This study presents a systems-level metabolic modelling framework for identifying tissue-specific collateral lethal vulnerabilities in cancer. By integrating genome-scale metabolic models with flux sampling and synthetic lethality analysis, we demonstrate that cancer-associated metabolic rewiring creates exploitable dependencies absent in normal tissue models. The absence of overlap in collateral lethal reactions across tissue types underscores the tissue specificity of these vulnerabilities and argues against the existence of universal metabolic targets, reinforcing the need for context-aware therapeutic strategies.

## 4. Methods

We begin by retrieving RNA-Seq transcriptomic data from TCGA for solid tumours, including samples from adjacent normal cells (Fig.1). This is followed by integration with genome-scale metabolic models to generate context-specific models for each tissue sample. FBA and flux sampling provide a mechanistic view of the metabolic capabilities of cancer tissues. Finally, using Fast-SL and DepMap, we identify and validate putative collateral-lethal genes.

### 4.1 Gene expression-guided reconstruction of context-specific metabolic models

We retrieved gene expression data from TCGA for eight tissues: breast, bronchus and lungs, colon, kidney, liver, prostate, stomach, and thyroid. Data from the GDC release 15.0 were used, with the selection criteria being “Primary Tumour” or “Solid Tissue Normal”. FPKM values were obtained for samples from both tumour and adjacent normal tissues. 30 random samples from each tissue were selected for further analysis.

A modified version of FASTCORE [10], an algorithm that reconstructs context-specific metabolic models from a global metabolic network, such as the Recon2.2 [39] GEM was used to reconstruct context-specific metabolic models for each tissue type in both cancer and normal conditions. Gene expression data were log-transformed, and genes were mapped to reactions using GPR (gene-protein-reaction) rules. Reactions were categorized into 4 categories based on the expression profiles: ZCR (zero-confidence reactions) with expression below the 10^th^ percentile, LCR (low-confidence reactions) with expression between the 10^th^ and 75^th^ percentile, MCR (medium-confidence reactions) with expression above the 75^th^ percentile, and HCR (high-confidence reactions) with expression above the 90^th^ percentile. The expressed reactions, along with the biomass and ATP-demand reactions, were used as a core reaction set to prune the generic Recon model with the FASTCORE algorithm. Penalties are assigned to reactions, with core reactions having lower costs and non-core reactions having higher costs. Optimisation minimises the penalties while ensuring model functionality. This is achieved iteratively, resulting in context-specific metabolic models constrained by sample-specific gene expression profiles and the metabolic reactions required for cell function. We generated 480 such metabolic models, 60 from each tissue type, with 30 models from each tumour and adjacent normal sample.

### 4.2 Outlier identification using DBSCAN

Pairwise metabolic distances (i.e., Jaccard distance) were computed between all context-specific models based on their reaction presence/absence profiles. Metabolic Distance = 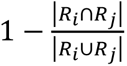, where *R_i_* is the reaction list from the model *i* and *R_j_* is the reaction list of the model *j*. A metabolic distance of 1 indicates that the two models do not share any reactions, whereas a metabolic distance of zero indicates that the models have identical reactions. Using this distance matrix, we performed density-based clustering to identify outliers using the *dbscan*, *fpc*, and *factoextra* packages in R. The *minPts* parameter was set to 3, and a k-nearest neighbour (kNN) distance plot was used to determine the optimal eps value. Out of 480 context-specific metabolic models, this clustering approach identified 85 outliers that were not restricted to any particular tissue type or condition. The remaining 395 models were retained for downstream analysis.

### 4.3 Differentially active pathways with flux sampling

We performed flux sampling using OptGpsampler^38^ in COBRApy [41]. 2000 samples were generated for each model. To account for baseline variations in total carbon influx and enable unbiased cross-tissue comparisons, flux values for all reactions were normalised to the glucose uptake rate of their respective models, and the mean relative flux was computed. Differential flux analysis between tumour and normal models was performed using the limma-eBayes framework. Pathway enrichment analysis of the differentially active reactions was carried out using the camera function in the limma [42] package. A logFC >= 1 and an FDR <= 0.05 were used as cutoffs for analysis.

### 4.4 Identification and validation of collateral-lethal gene pairs with Fast-SL and DepMap

Fast-SL[12] predicts essential genes and reactions, called ‘synthetic lethals’ in metabolic networks. The method predicts single lethal (SL) and higher-order lethal events, such as double lethal (DL) and triple lethal. We utilised Fast-SL [12] to identify single- and double-lethal reactions across all cancer and normal tissue metabolic models. The objective function was set to maximise biomass production to identify synthetic lethal reactions. We used Fisher’s exact test to determine statistically significant single lethal reactions in cancer and double-lethal reaction pairs in normal models. SL reactions in cancer models that were a component of DL reaction pairs of the normal models were classified as ‘collateral lethal’. Details of the list of lethal genes identified are provided in Supplementary Table S2.

To validate the predicted collateral lethal reactions, we used the DepMap [43] 22Q2 version in R using the ‘*depmap*’ package, which contains comprehensive cancer dependency data, including gene essentiality across various cell lines. We use CRISPR data, gene mutations, and copy-number details to evaluate the predicted collateral lethal pair with greater confidence. For a gene A-gene B pair, we first identify cell lines where gene B is mutated and check if gene A is essential and expressed in gene B-inactivated cell lines. Gene pairs were classified as ‘valid’ collateral lethal interactions if gene A was essential and expressed in at least 30% of gene B-inactivated cell lines (Supplementary Table S3). This threshold was empirically established to capture biologically significant, recurrent collateral dependencies. We further utilised the Online Gene Essentiality [44] (OGEE) Database (https://v3.ogee.info) to reach a consensus on the predicted collateral lethality.

### 4.5 Metabolic rewiring in cancer models based on flux distributions

For the collateral lethal gene pairs identified as ‘valid’, we deleted gene B from the models representing the normal tissue state and compared the FBA flux distributions for all reactions with those in the corresponding cancer models. Theoretically, deletion of gene B in normal tissue models is expected to induce compensatory metabolic adaptations that may partially recapitulate flux redistribution patterns observed in the corresponding cancer models. Deviations in flux distributions between perturbed normal and cancer models may therefore highlight additional cancer-specific metabolic rewiring associated with oncogenic transformation. Any significant changes in flux patterns among shared reactions are noted. Reactions that are active (i.e., fluxes change from zero to an absolute flux value of 0.01 or higher) or inactive (i.e., fluxes change from an absolute flux value of 0.01 or higher to zero) between the cancer and normal models highlight the metabolic rewiring that occurs in cancer cells.

## Code availability

Code repository for the data analysis can be found at https://github.com/RamanLab/cancer_collaterallethality

## Acknowledgements

The results presented here are partly based on data from the TCGA Research Network (https://www.cancer.gov/tcga).

M.I. and R.B. acknowledge research fellowships from the Centre for Integrative Biology and Systems mEdicine, Indian Institute of Technology, Madras.

## Author Contributions

M.L. and K.R. conceived the original study. M.I. and R.B. conducted the data analysis. M.I. and M.L. wrote and revised the paper with comments from K.R.

## Supplementary Figures

**Supplementary Fig. S1.**
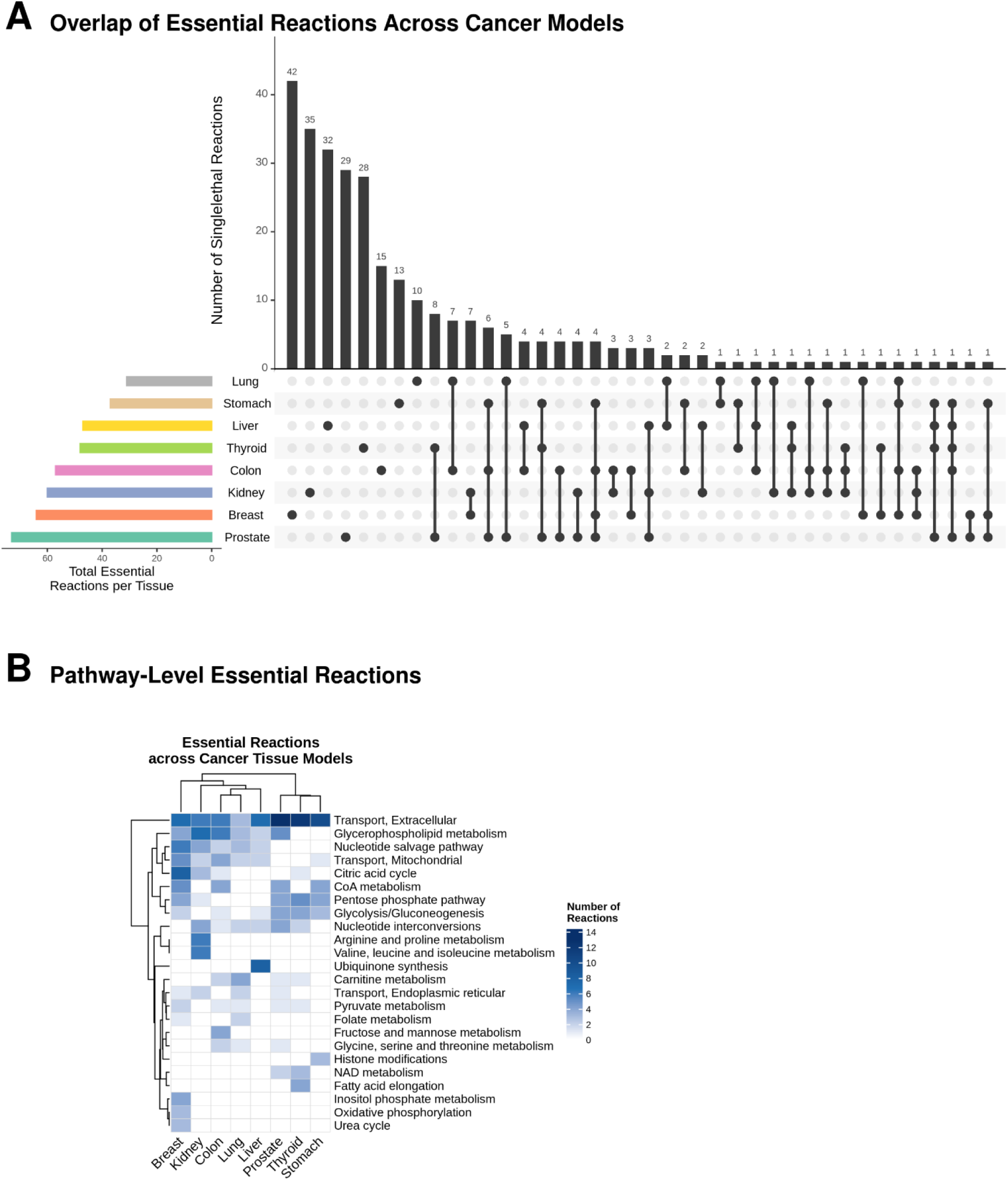
**S1A:** Overlap of essential reactions across tumour-specific metabolic models. **S1B:** Metabolic subsystems associated with the essential reactions, highlighting similarities in essentiality profiles among different tissue types.

**Supplementary Fig. S2.**
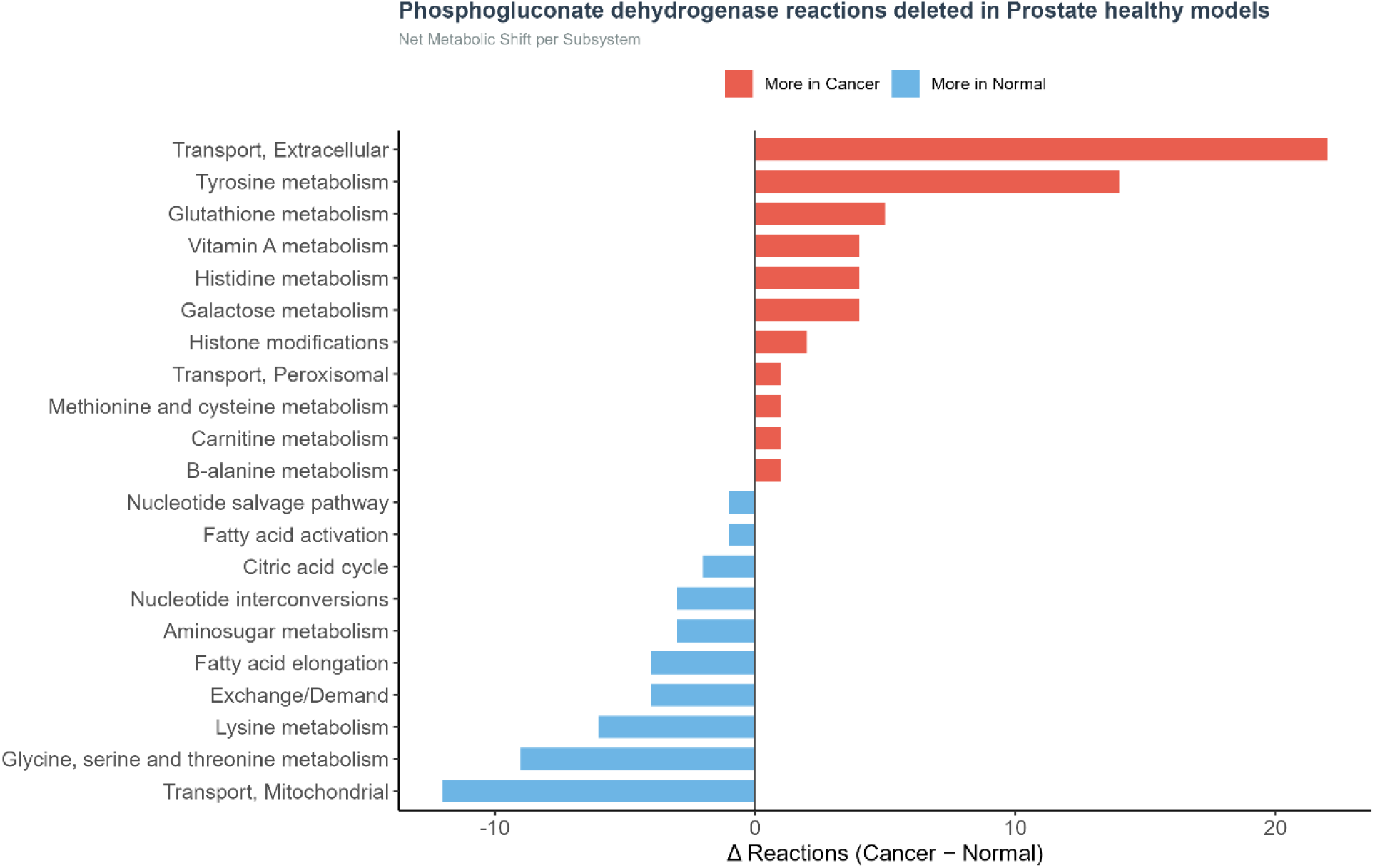
The phosphogluconate dehydrogenase reaction was deleted from normal prostate tissue metabolic models, and the resulting metabolic rewiring was evaluated by comparing the number of reactions within subsystems that showed higher activity in cancer versus normal models.

**Supplementary Fig. S3.**
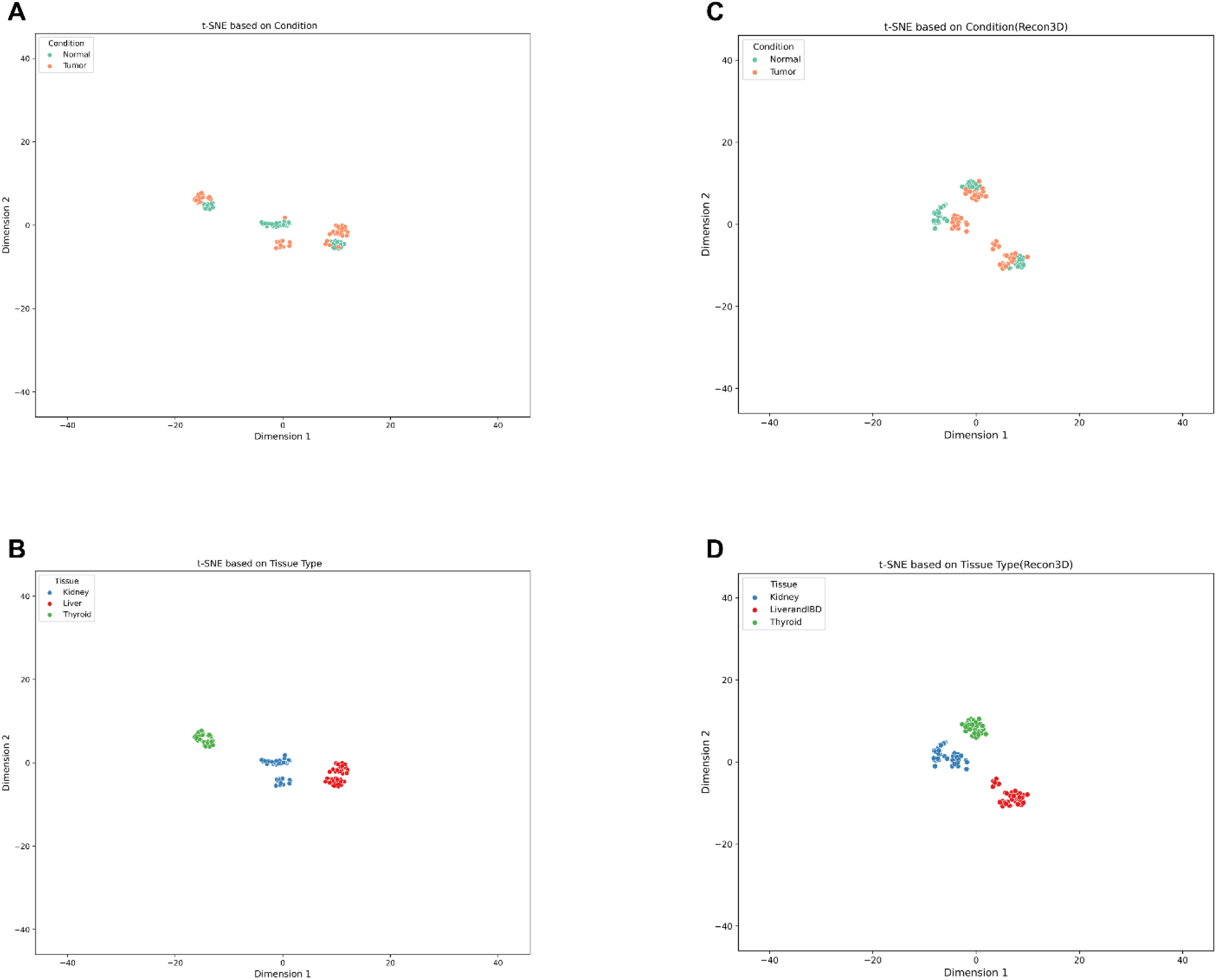
Two-dimensional t-SNE visualisations comparing models generated from the modified Recon2.2 reference GEM used for all primary analyses in this study (A and B) against models generated from the Recon3D reference GEM (C and D). Panels A and C illustrate clustering by condition (Normal vs. Tumour), while panels B and D illustrate clustering by tissue type.

